# Epigenomic and functional dynamics of human bone marrow myeloid differentiation to mature blood neutrophils

**DOI:** 10.1101/295014

**Authors:** Luigi Grassi, Farzin Pourfarzad, Sebastian Ullrich, Angelika Merkel, Felipe Were, Enrique Carrillo de Santa Pau, Guoqiang Yi, Ida H Hiemstra, Anton TJ Tool, Erik Mul, Juliane Perner, Eva Janssen-Megens, Kim Berentsen, Hinri Kerstens, Ehsan Habibi, Marta Gut, Marie Laure Yaspo, Matthias Linser, Ernesto Lowy, Avik Datta, Laura Clarke, Paul Flicek, Martin Vingron, Dirk Roos, Timo K van den Berg, Simon Heath, Daniel Rico, Mattia Frontini, Myrto Kostadima, Ivo Gut, Alfonso Valencia, Willem H Ouwehand, Hendrik G Stunnenberg, Joost HA Martens, Taco W Kuijpers

## Abstract

Neutrophils are short-lived blood cells that play a critical role in host defense against infections. To better comprehend neutrophil functions and their regulation, we provide a complete epigenetic and functional overview of their differentiation stages from bone marrow-residing progenitors to mature circulating cells. Integration of epigenetic and transcriptome dynamics reveals an enforced regulation of differentiation, through cellular functions such as: release of proteases, respiratory burst, cell cycle regulation and apoptosis. We observe an early establishment of the cytotoxic capability, whilst the signaling components that activate antimicrobial mechanisms are transcribed at later stages, outside the bone marrow, thus preventing toxic effects in the bone marrow niche. Altogether, these data reveal how the developmental dynamics of the epigenetic landscape orchestrate the daily production of large number of neutrophils required for innate host defense and provide a comprehensive overview of the epigenomes of differentiating human neutrophils.

**Key points:**

- Dynamic acetylation enforces human neutrophil progenitor differentiation.

- Neutrophils cytotoxic capability is established early at the (pro)myelocyte stage.

- Coordinated signaling component expression prevents unwanted toxic effects to the bone marrow niche.

## Introduction

Human neutrophils, with a daily production rate of ~10^11^ cells (Strydom and Rankin, 2013), are the most abundant type of leukocytes and an essential element of the innate immune system. Patients with either acquired or inherited defects in neutrophil development, migration or function show enhanced susceptibility to infection by opportunistic pathogens (Erlacher and Strahm, 2015, Fodil et al., 2016). Neutrophils have a variety of effector functions, such as phagocytosis of pathogens (Nauseef and Borregaard, 2014), intracellular killing in phagosomes with reactive oxygen species (ROS), and anti-microbial granule components (Darrah and Andrade, 2012, Garcia-Romo et al., 2011, Hakkim et al., 2010). They are produced in the bone marrow and reach the bloodstream across five main stages of differentiation. Myeloblasts (MBs) undergo one week of proliferation followed by maturation (Bainton et al., 1971), then progressively move through (pro)myelocytes (P/Ms), metamyelocytes (MMs), band neutrophils (BNs) and finally segmented neutrophils (SNs). During this time they produce different granules filled with antimicrobial compounds and develop the abilities of respiratory burst and chemotaxis. SNs represent a reserve pool from which cells are constantly released into the peripheral bloodstream, as highly toxic non–dividing polymorphonuclear neutrophils (PMNs). These cells have a short circulating half-life of five to eight hours before they extravasate to patrol tissues (Galli et al., 2011). The differentiation process exhibits a precise and tightly defined transcriptional program (Theilgaard-Monch et al., 2005). In this study, we used total RNA-sequencing (RNA-seq) to characterize and compare gene expression profiles of the various differentiation stages. Histone modifications, chromatin immunoprecipitation sequencing (ChIP-seq) and whole-genome-bisulfite-sequencing (WGBS) have been used to describe the epigenetic landscapes underlying these coordinated changes. For each stage we defined DNA methylation patterns and characterized four active histone marks: H3 lysine-4 trimethylation (H3K4me3), H3 lysine-27 acetylation (H3K27ac), H3 lysine-4 monomethylation (H3K4me1), H3 lysine-36 trimethylation (H3K36me3) and two repressive marks: histone H3 lysine-9 trimethylation (H3K9me3) and histone H3 lysine-27 trimethylation (H3K27me3). The integration of these analyses reveals a high degree of coordination between the dynamics of histone modifications and transcriptional changes. The activation of PMNs triggers the host defense. This process is accomplished through some functional key responses as respiratory burst, cell migration, and degranulation. Integrating our functional assays with the epigenomic analysis gave us the opportunity to further characterize the timing of these important processes.

## Results

### Transcriptional changes in neutrophil development

To better understand neutrophil functions and their regulation we analyzed the transcriptomes of circulating neutrophils and their progenitor populations (supplementary Figure 1A). Our analysis revealed quantitative and qualitative differences in expression between PMNs, SNs and the other differentiation stages. We used cumulative distribution of expression to explore the complexity of the transcriptome (Mele et al., 2015) and found that in PMNs and SNs, the genes with higher expression give a smaller contribution to the global transcriptional output than in other stages (Figure 1A). To deepen the transcriptome comparisons we performed differential gene expression analysis between subsequent stages, and as well, between the initial and the final stage (Figure 1B and supplementary data 1). Down-regulated genes in the P/M-PMNs comparison enriched gene ontology (GO) categories related to cell functions such as metabolism, translation, cell cycle, DNA replication, mitochondria and cell divisions (supplementary Figures 1B-1C). On the other hand, the up-regulated genes enriched fewer categories, mainly related to immune system functions and vesicle transport processes (supplementary Figures 1B-1C). Cluster analysis and gap statistic (Tibshirani et al., 2001) summarized the differentially expressed genes in at least one of the five comparisons (P/M-MM, MM-BN, BN-SN, SN-PMN, P/M-PMN) in 7 main clusters (r1 to r7, Figure 1C and supplementary data 1). Genes in clusters r1 and r2 showed an increase in expression toward the latter stages of differentiation and enriched GO terms related to immune system and signal transduction (supplementary data 1). Genes in clusters r5, r6 and r7 displayed a decreasing expression trend with differentiation progression and enriched GO terms related to translation, mitochondria and cell cycle (supplementary data 1). The remaining two clusters (r3 and r4) were made by genes whose expression increased during the intermediate stages of differentiation and then decreased at PMN stage. These enriched GO terms related to vesicle transport and signal transduction (supplementary data 1).

**Figure 1.**
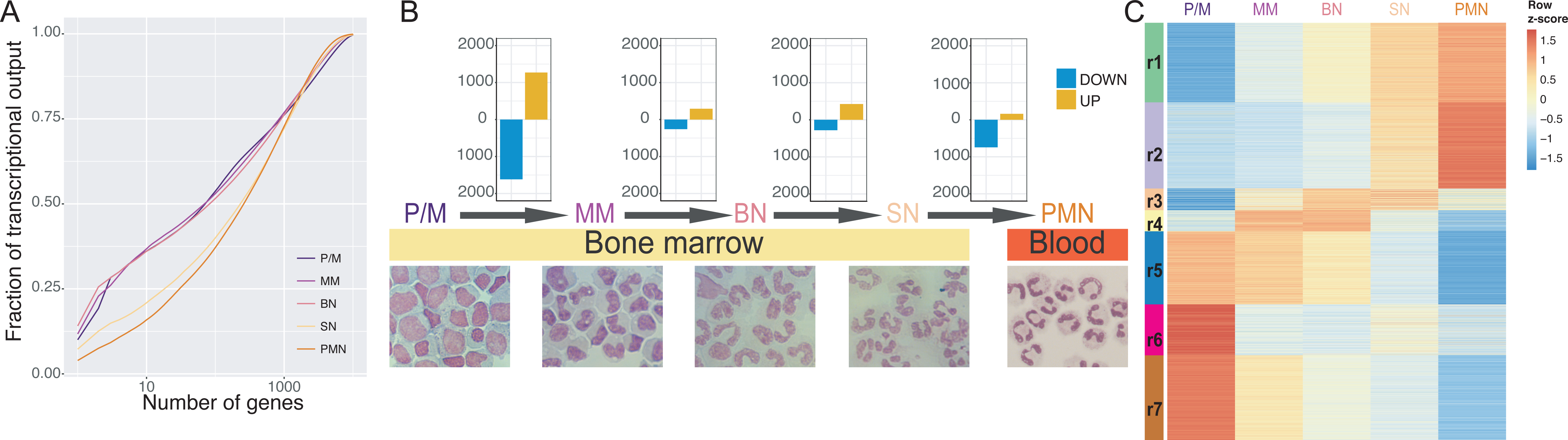
Transcriptome and epigenome dynamics of neutrophil differentiation. **(A)** Cumulative distribution of the average fraction of total transcription contributed by genes when sorted from most-to-least expressed in each differentiation stage. **(B)** Top panel: Bar plots reporting differentially expressed genes (posterior probability > 0.5 and absolute fold change > 2) in the comparisons P/M-MM, MM-BN, BN-SN, SN-PMN. Bottom panel: May-Grunwald-Giemsa staining of the isolated cells. **(C)** Heatmap displaying the expression patterns of the clusters (r1-r7) identified by the K-means analysis of the genes differentially expressed in at least one comparison.

### Epigenetic dynamics in neutrophil development

In order to define the chromatin dynamics underlying the changes in gene expression during neutrophil differentiation we determined the epigenetic landscape in the five differentiation stages and integrated them with the results of the transcriptome characterization. Principal component analysis (PCA) of the four active histone modifications (H3K4me3, H3K27ac, H3K4me1 and H3K36me3) revealed a good separation among the five neutrophil development stages (supplementary Figure 2A). In contrast, PCA for the two repressive marks (H3K9me3 and H3K27me3) revealed donor-specific, rather than stage specific, clusters (supplementary Figure 2A). Moreover, PCA analysis of DNA methylation profiles did not reveal differences among the five differentiation stages (supplementary Figure 2B) and only few differentially methylated regions (DMRs) were identified between subsequent stages of differentiation (supplementary Figure 2C and supplementary data 2). Altogether histone mark and DNA methylation results indicate a modest contribution of repressive epigenetic mechanisms in the described phases of neutrophil differentiation.

**Figure 2.**
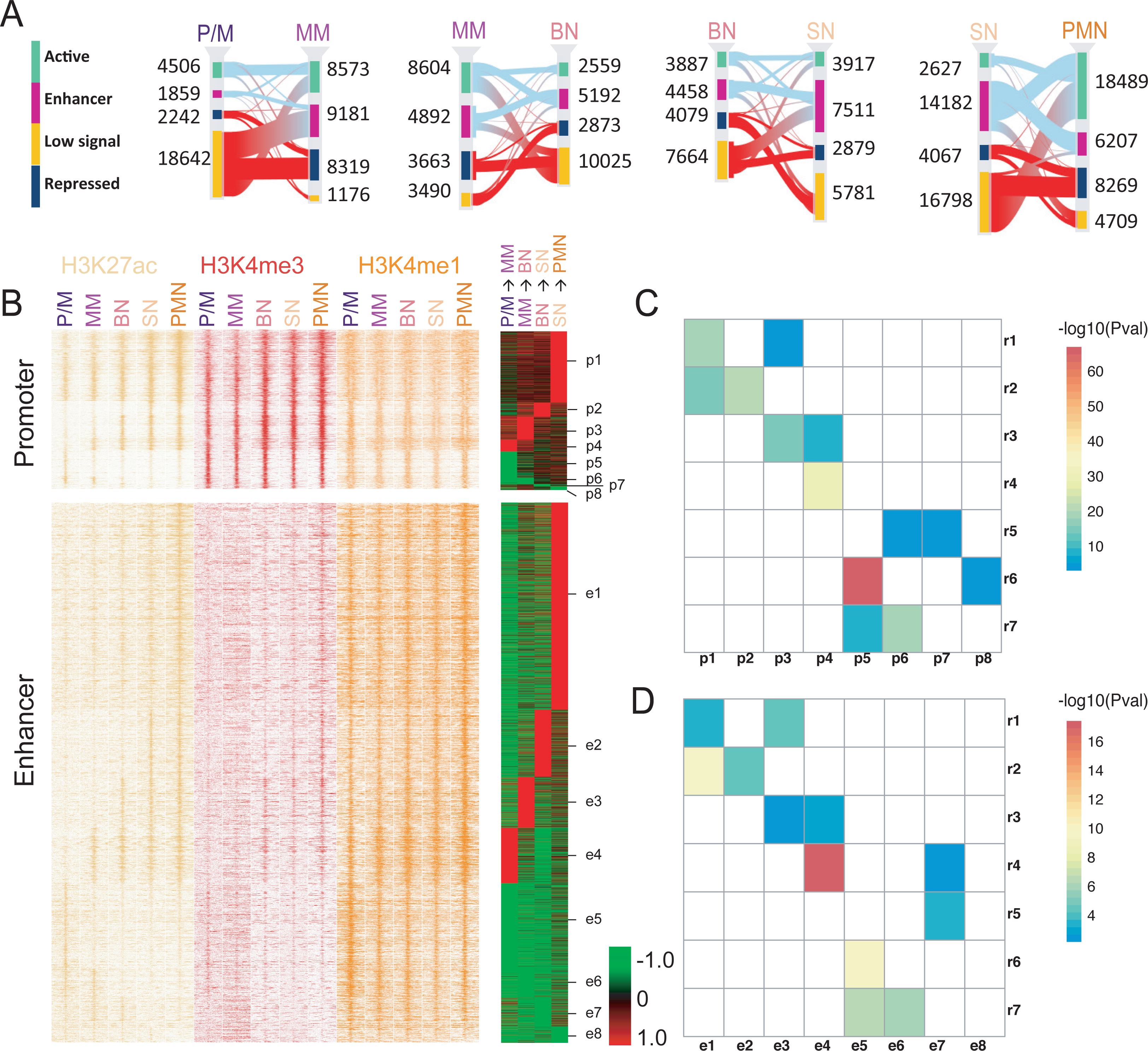
H3K27 acetylation drives consecutive neutrophil differentiation stages. **(A)** Sankey diagrams of chromatin state changes for each of the neutrophil differentiation transitions. Different chromatin states are represented with specific color codes: green indicates active regions, purple enhancers, amber low signal regions and dark blue heterochromatic regions. The red connecting lines indicate transitions towards inactive states (low signal and repressed); the light blue lines indicate transitions towards active states (active and enhancer). **(B)** Left panels, Stacking heat maps of H3K27ac (yellow), H3K4me3 (red) and H3K4me1 (orange) ChIP-seq of promoters and enhancers (± 5 kb) with dynamic acetylation during different stages of neutrophil differentiation. Right panels, differentially acetylated promoter and enhancer during two consecutive neutrophil progenitor stage transitions are clustered as acetylation clusters for promoters p1 to p8 and for enhancers e1 to e8, respectively. Acetylation gains are presented in red and acetylation losses in green. **(C-D)** Heatmaps representing the association between the expression clusters identified in figure 1C and the promoter and enhancer clusters identified in figure 2B. Only the significant corrected *P* values (<0.05) are reported in the figures.

The six different histone modifications were also used to perform a genome segmentation analysis (Ernst and Kellis, 2012) which indicated 12 representative states (supplementary Figure 3A). These were collapsed into a smaller set for ease of interpretation (Carrillo-de-Santa-Pau et al., 2017) to obtain 4 main categories: active, enhancer, low signal and repressed (supplementary Figure 3A). We then focused on the changes of chromatin states, identifying between 15,685 and 27,604 at each consecutive differentiation transition (Figure 2A and supplementary data 3). In agreement with the differential gene expression analysis results, the consecutive transitions with most changes were P/M to MM and SN to PMN. Up-regulated genes were enriched in changes towards active states (i.e. from low signal/repressed to active/enhancer), whilst down-regulated genes were enriched in transitions towards inactive states (i.e. from active/enhancer to low signal/repressed) in the majority of differentiation stages (supplementary Figure 3B). H3K27ac, strongly associated with active and enhancer states, is among the most highly informative marks about gene regulation (Karlic et al., 2010, Rada-Iglesias et al., 2011). For this reason, we decided to focus our attention on the 9100 genomic regions characterized by changes in H3K27ac levels during differentiation (see Experimental procedures). Using co-occupancy with H3K4me3 to identify promoter regions and co-occupancy with H3K4me1 to identify enhancers, in addition to a supervised cluster of the H3K27ac log ratio rpkm, we divided these regions in two categories each made of 8 subgroups (Figure 2B and supplementary data 4). The intersection of these regions with the chromatin states revealed an overlap with active (promoter) regions in the H3K4me3-enriched clusters and an overlap with enhancers in the H3K4me1-enriched clusters (supplementary Figure 3C), corroborating our partitioning. We refer to the clusters as promoter (p1-p8) and enhancer (e1-e8). In both promoter and enhancer clusters ~2/3 of dynamic regions displayed an increase in H3K27ac (i.e. increase in activity), whilst ~1/3 displayed a reduction in H3K27ac (i.e. reduction in activity). The contingency tables made by the intersection of the genes in the RNA-seq clusters gave significant Pearson’s chi-squared for both acetylation clusters (enhancer chi-squared = 513, *P* value = 0.0005; promoter chi-squared = 1364, *P* value = 0.0005). We tested the association of each gene expression cluster with each dynamic acetylation cluster (Figures 2C and 2D). Interestingly we found that expression clusters r1 and r2, containing genes with increased expression, were significantly associated with promoter clusters p1 and p2 and enhancer clusters e1 and e2. Expression clusters r6 and r7, formed by genes with decreased expression, were significantly associated with promoter clusters p5, p6 and enhancer clusters e5 and e6, all characterized by reduced histone acetylation levels. The expression cluster r4 included genes with an increased expression in the PM to MM transition and the same trend was observed in the associated acetylation clusters e4/p4.

**Figure 3.**
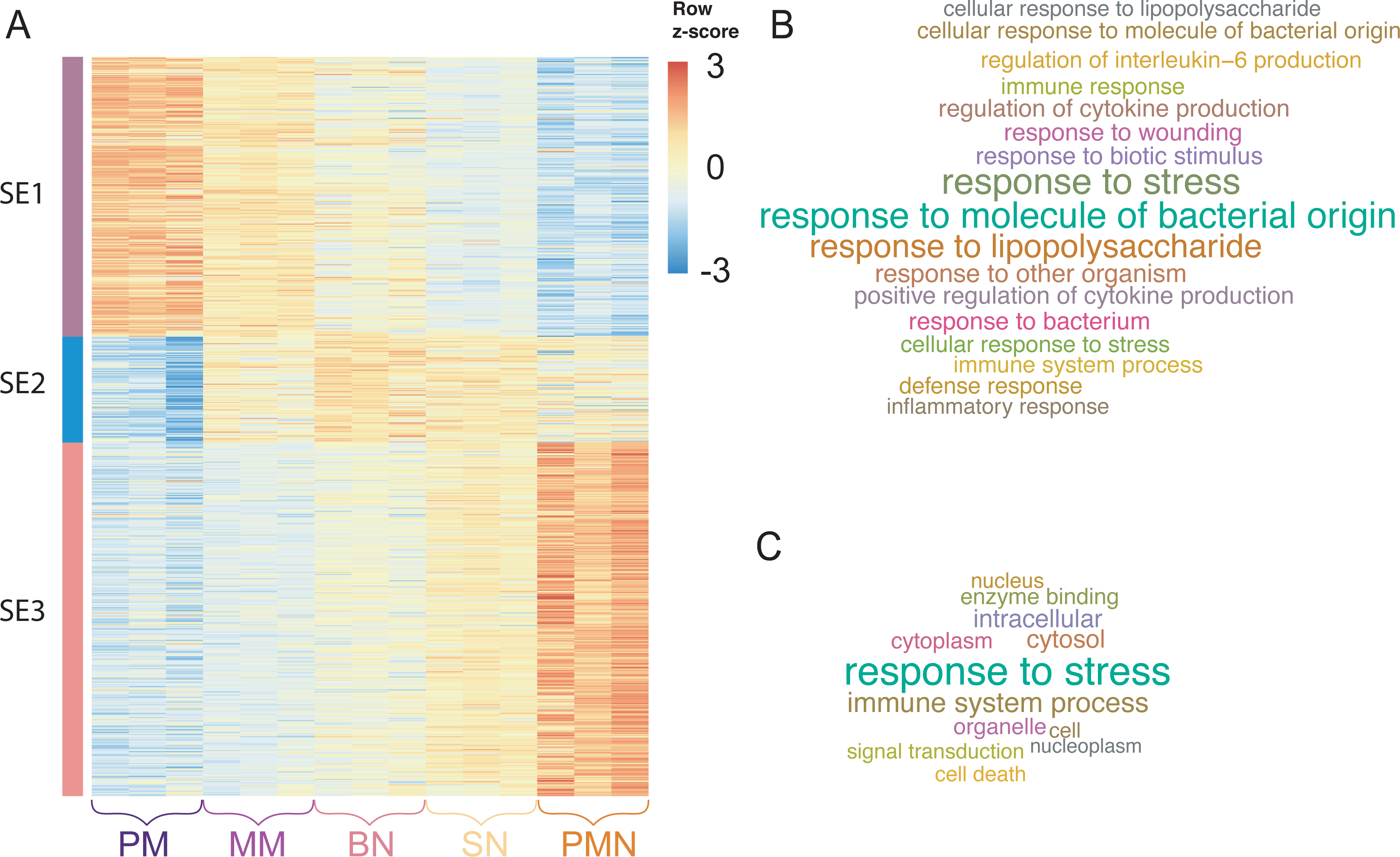
Identification of super-enhancers associated with neutrophil differentiation. **(A)**Heatmap of H3K27ac density at super-enhancers with dynamic acetylation during different stages of neutrophil differentiation. For each state tag density for the three replicates is included. **(B**) GO Biological Processes and GO Slim **(C)** categories enriched by genes associated with super-enhancers activated at the final stages of neutrophil differentiation (cluster SE3).

Dynamically acetylated enhancers of the same cluster had a genomic distance closer than expected by chance (permutation test z = 21). This observation prompted us to investigate the presence of super-enhancers (SEs), regulatory elements made by clusters of enhancer that specify cell identity (Whyte et al., 2013, Witte et al., 2015). We made use of the established ROSE algorithm to identify 629 SEs, which could be clustered in 3 main groups (Figure 3A and supplementary data 4). The SE cluster 1 (SE1) was made by elements more acetylated in the P/M state, showing a decreasing trend of acetylation in the subsequent stages differentiation, the cluster SE2 had an opposite trend, with the acetylation increasing in the first stages and then decreasing at the SN-PMN transition. Finally, cluster SE3 contained SEs activated at the end of differentiation process. Genes controlled by SEs belonging to cluster 3 were the only ones showing an enrichment for functional categories, all of them related to PMN’s activation (Figure 3B-C).

### Transcription factors and release from bone marrow

Transcriptional programs are enforced at epigenetic level by transcription factors (TFs) binding at regulatory elements (promoters and enhancers) (Lambert et al., 2018, White et al., 2013). To identify the transcription factors that have an important role in granulopoiesis we searched for TF family consensus binding sites, enriched in regions with increased acetylation (clusters p1-p4 and e1-e4) compared to regions with decreased acetylation (clusters p5-p8 and e5-e8) and *vice versa.* We found enrichments for basic leucine zippers (bZIP), SOX, zinc fingers (ZF), nuclear receptors (NR), basic helix-loop-helices (bHLH), NF-kB family members (REL), homeodomains and ETS families, with stronger signal in regions with increased acetylation (Figure 4A). Regions with decreased H3K27ac signal showed enrichment for MYB- and GATA-binding motifs. We integrated these results with gene expression analysis results and identified for each enriched motif TF genes differentially expressed in at least one of the comparisons between differentiation stages (Figure 4B). The majority of these TFs (81 of 95) were differentially expressed in only one of the four comparisons. Twelve of the remaining fourteen genes, differentially expressed at multiple transitions, were always either up or down-regulated. Only two genes, differentially expressed at multiple transitions had a trend inversion: TGIF2 was down-regulated in the P/M-MM transition and up-regulated in the BN-SN transition and ETS1 up-regulated in the P/M-MM transition and down-regulated in the SN-PMN transition.

**Figure 4.**
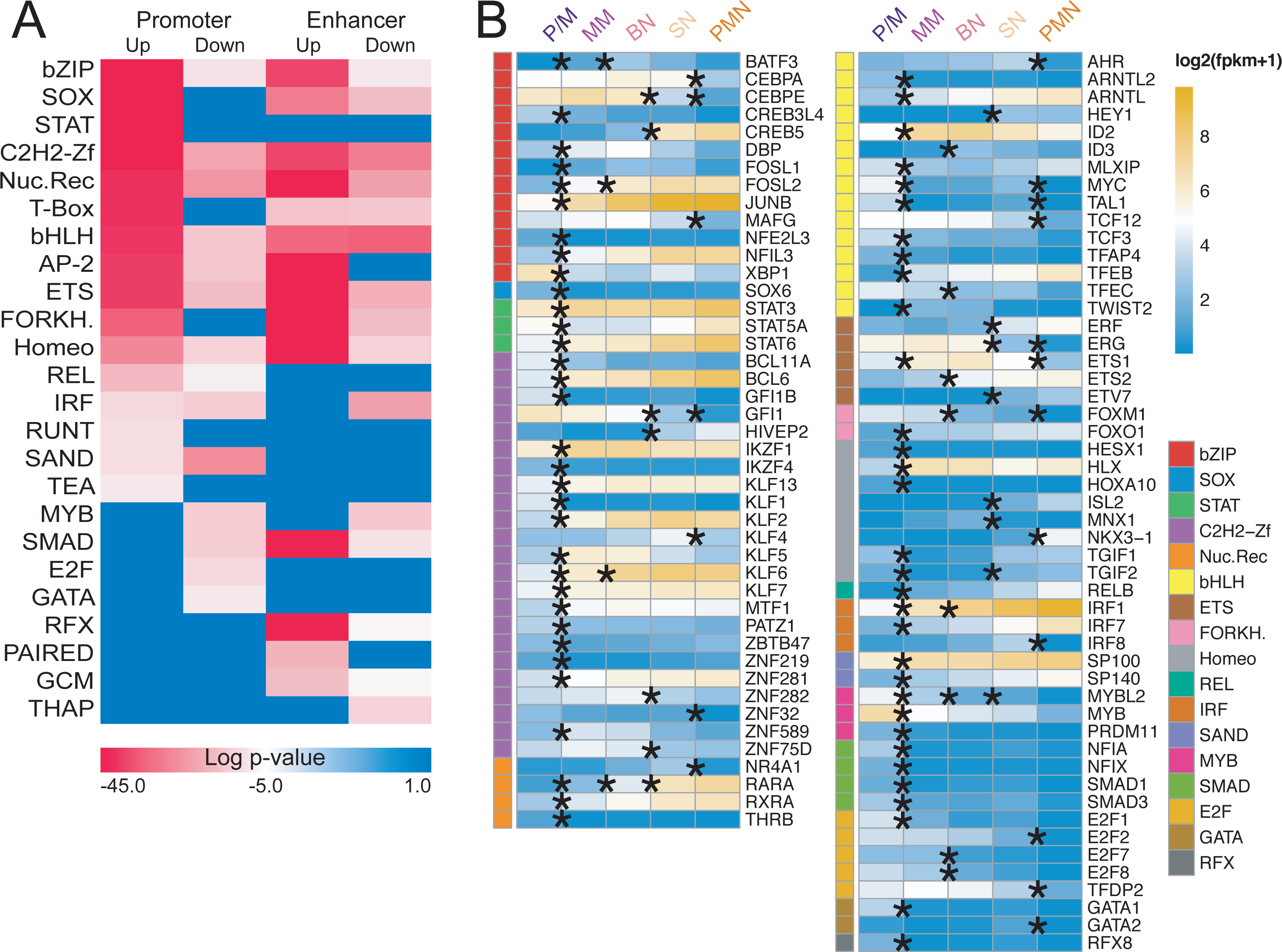
Transcription factors with enriched binding sites in dynamic acetylated regions. **(A)** Transcription factor (TF) family motif enrichment in the dynamic acetylated regions in figure 2B. **(B)** RNA expression in log2(FPKM+1) of the TF family members in (A) differentially expressed in at least one of the comparison between differentiation stages, the * denotes the transitions where the expression is significantly different.

The P/M to MM transition included the majority (65 of 95) of differentially expressed TFs followed by the bone marrow-to-blood transition (SN to PMN) with 20 differentially expressed TFs (Figure 4B). Interestingly 19 of them were down-regulated and NKX3-1, the only up-regulated one, had a reported transcriptional repressor activity (Simmons and Horowitz, 2006). The TF families MYB, GATA and E2F were exclusively overrepresented in enhancers/promoters with decreased acetylation, reflecting their role in other lineages and cell cycle progression (Bartunek et al., 2003, Ghazaryan et al., 2014). Accordingly, differentially expressed genes from these families were all down-regulated (Figure 4B).

### Acquisition and control of cytotoxic capabilities

Cell proliferation is halted during neutrophil differentiation; our transcriptome and epigenome data indicated that this occurs in the bone marrow, as previously reported (Klausen et al., 2004, Theilgaard-Monch et al., 2005). Flow cytometry quantification of Ki67, a marker for proliferation (Gerdes, 1990), showed that the majority of the signal drops in the early stages of myeloid differentiation (P/M-to-MM transition) and a complete loss is observed at the SN stage (Figure 5A). Neutrophils are phagocytes, capable of ingesting and killing microorganisms by the release of ROS, highly toxic chemical species, potentially harmful for the bone marrow niche. The multicomponent nicotinamide adenine dinucleotide phosphate (NADPH) oxidase complex, consists of a membrane-associated heterodimer cytochrome *b*_558_ and a trio of cytosolic proteins (Nauseef and Borregaard, 2014). This complex confers the unique neutrophils’ ability to generate a family of reactive oxidizing (Weiss, 1989) and we focused on it to characterize the exact dynamic of this defense mechanism formation. Upon activation, the intact NADPH oxidase generates superoxide, the precursor to hydrogen peroxide and other ROS with microbial activity, at the plasma membrane and at the membranes of phagosomes where particles are ingested (Figure 5B). The oxidase complex subunits showed different expression dynamics during differentiation (Figure 5C). CYBA (p22-phox), NCF4 (p40-phox), RAC2 and RAC1 had stable expression levels, whereas CYBB (gp91-phox) was maximally expressed between the MM and BN stages and the cytoplasmic components NCF1 (p47-phox) and NCF2 (p67-phox) were up-regulated in the P/M to MM transition and then remained stable in the subsequent stages. In order to verify when the complex is functional, we used an Amplex Red hydrogen peroxide assay to detect the NADPH oxidase activity after the supraphysiological stimulation of PAF-primed neutrophils with the bacterial-derived tripeptide formyl-MLP (PAF/fMLP), a compound able to activate the complex above natural stimulation (Drewniak et al., 2013, Kuijpers et al., 2005). At P/M the NAPDH oxidase activity was very low. It became readily detectable at the next stage of differentiation (MM) as shown by the activation with the cell-permeable phorbol ester PMA, which directly activates intracellular protein kinase C (Moore et al., 1991). In contrast, stimulation via the cell-impermeable PAF/fMLP revealed a delay in ROS formation, detectable from the BN stage onwards (Figure 5D). In agreement with the increase in transcription level and functional activity of NADPH oxidase we found that cytochrome b_558_, the heterodimer of gp91-phox and p22-phox, and fMLP-receptor FPR1 levels (both determined by flow cytometry) increased during differentiation (supplementary Figures 4A and 4B). Additionally we also noted that CXCR1, CXCR2, CXCR4, FPR1, FPR2, PTAFR and C5AR1, activating G-protein-coupled receptors for the major chemotactic factors in neutrophils, follow the same expression trend (supplementary Figure 4C) and are all in the expression cluster r1.

**Figure 5.**
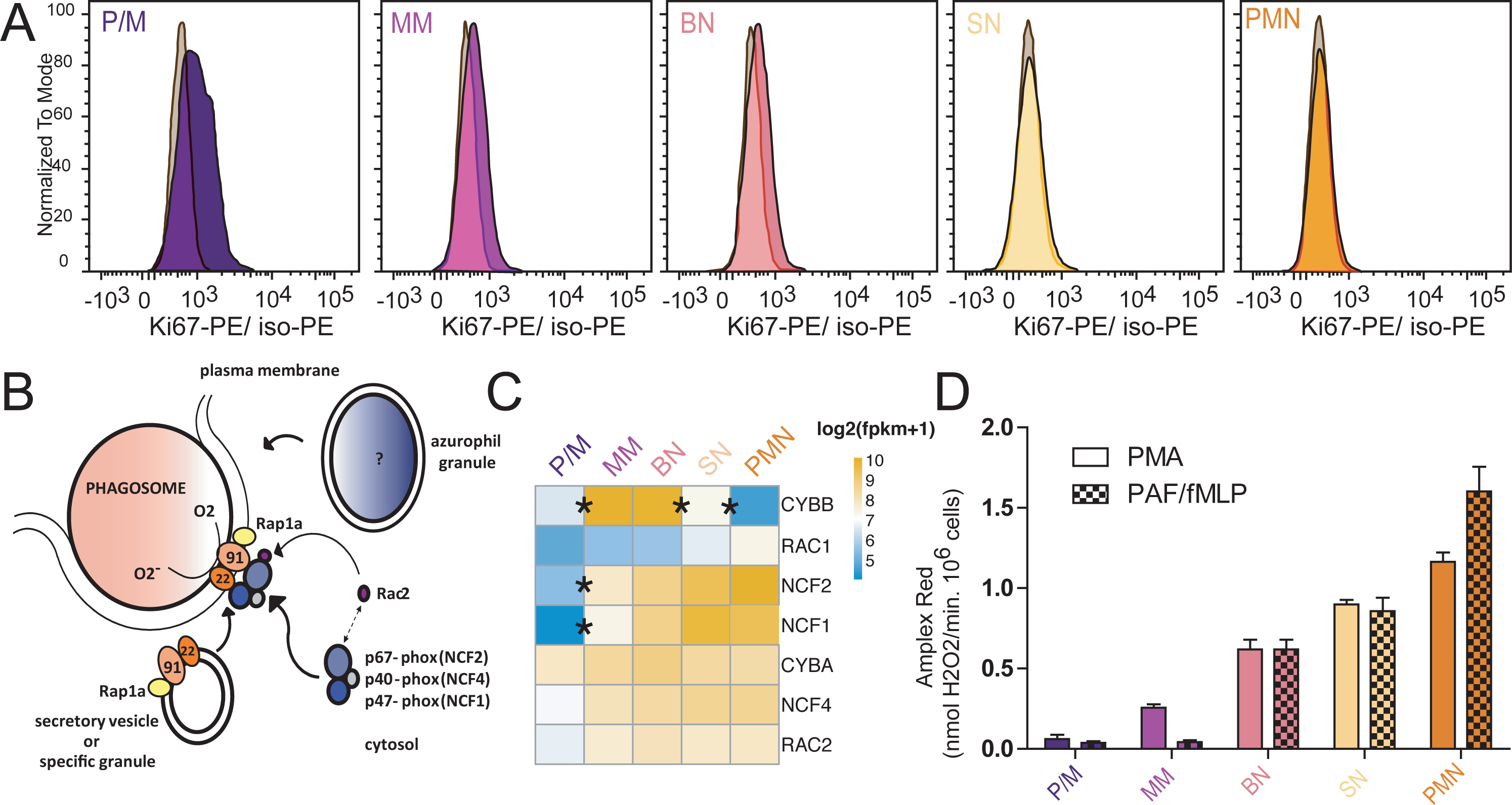
Cell cycle and NADPH oxidase regulation in neutrophil differentiation. **(A)** Flow cytometry analysis of Ki67 expression in bone marrow neutrophil progenitor fractions normalized to isotope control in grey (n=3). **(B)** Schematic of the assembly of the nicotinamide adenine dinucleotide phosphate (NADPH) oxidase enzyme complex on the phagosomal or plasma membrane upon neutrophil activation. **(C)** Heatmap representing the expression in log2(FPKM+1) of NADPH complex subunits during neutrophil differentiation. **(D)** Respiratory burst in sorted neutrophil progenitors from bone marrow and mature PMN upon stimulation with PMA and PAF/fMLP. Results are expressed as maximal rates in nmol H_2_O_2_/min.10^6^ PMNs, and as mean ± SEM (n=3). T-test *P* values of the different comparisons are reported in supplementary data 5.

### Transcriptional and epigenetic control of granule proteins

Another main function of the neutrophils is the degranulation consisting in the release of an assortment of proteins, stored in different granules. These granules are distinguishable by their contents: azurophilic granules (AG), specific granules (SG), gelatinase-containing granules (GG), ficolin-containing granules (FG), and secretory vesicles (SV) (Rorvig et al., 2013). To gain insight in the control of the biogenesis of these granules during differentiation, we examined available proteomic data (Rorvig et al., 2013) and selected those proteins whose granule assignments were supported by other existing literature (supplementary data 6). We found that genes encoding proteins belonging to various granules display different expression trends (Figure 6A), and, in agreement with this, they showed a tendency to belong to different RNA-Seq clusters (chi-squared = 91.8062, *P* value < 0.0005, supplementary figure 5A). AG proteins were associated with clusters r6 and r7 (Fisher exact test corrected *P* value < 0.05), whilst CM proteins were associated with cluster r1 (Fisher exact test corrected *P* value < 0.005) and SG proteins with cluster r4 (Fisher exact test corrected *P* value = 3.1 x 10^-6^).

**Figure 6.**
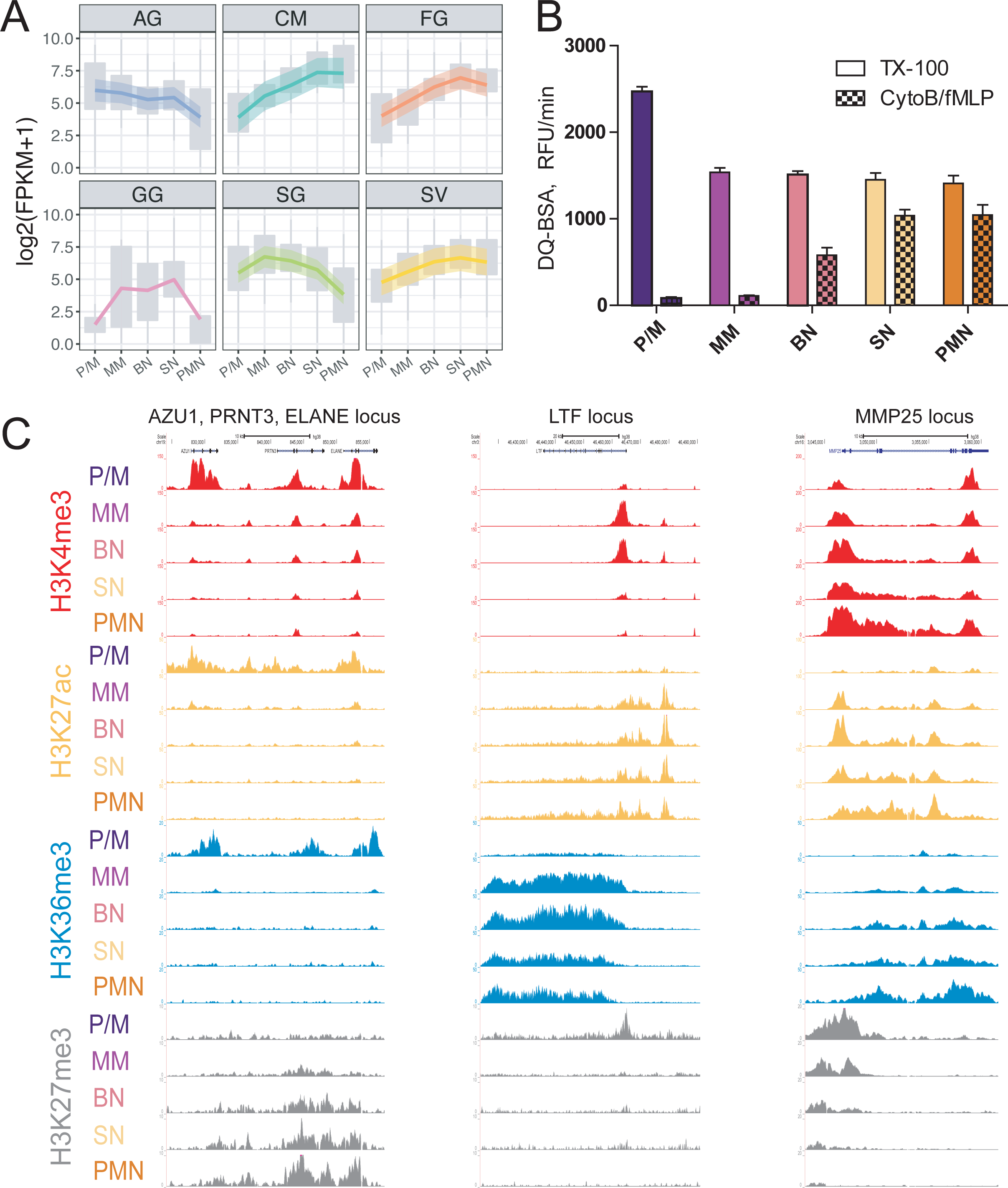
Epigenetics and transcriptomics of granule proteins. **(A)** Granule protein expression across differentiation. Upper and lower box plot margins indicate first and third quartile. LOESS fitting of the data with relative confidence interval is represented by a colored line with a shadow area **(B)** Proteolytic activity of the different differentiation stages measured by DQ-BSA cleavage. Triton X-100 releases total proteolytic activity stored in the cell by lysis, while proteolytic activity by CytoB/fMLP is a receptor-mediated release of proteases. Results are shown as the maximal slopes in RFU/min and expressed as mean ± SEM (n=3). T-test *P* values relative to the different comparisons are reported in supplementary data 5. **(C)** Representative views of epigenetic modifications dynamics at 3 loci encoding for granule proteins.

To determine the time of appearance of azurophilic granule’s proteolytic capability, we measured their combined serine protease activity, in sorted neutrophil progenitors, upon CytoB/fMLP stimulation or Triton X-100 lysis. The results showed that, while AG proteins have already been produced and stored at P/M stage, the protease release machinery is not in place before the BN stage. P/M cells were found to harbor the highest concentration of proteolytic activity, which was then diluted over to the MM daughter cells and remained stable in activity thereafter (Figure 6B). We also associated granule gene expression with the H3K27ac dynamic clusters, finding that genes encoding soluble AG proteins, such as *AZU1*, *PRTN3* and *ELANE* were enriched in clusters p5, p6 and e5 (corrected *P* values of the Fisher test: p5 = 6.64 x 10^-3^, p6 = 6.64 x 10^-3^, e5 = 3.65 x 10^-2^), with H3K27ac enriched in the P/M stage and reduced in the MM stage, as exemplified in Figure 6C. In contrast, SG-associated genes, such as *LTF*, were mainly enriched in clusters p4 and e4 (corrected *P* values of the Fisher test: p4 = 1.7 x 10^-3^, e4 = 2.88 x 10^-2^), representing increased acetylation in the P/M to MM transition.

## Discussion

Our study dissects human neutrophil differentiation through the analysis and the integration of transcriptomes, epigenomes and functional assays of five consecutive stages of maturation.

In the transcriptome of a cell a relative small number of genes contribute to a large fraction of the expression (Mele et al., 2015). Interestingly, we found this trend to be less pronounced in the two latest stages of neutrophil maturation, SN and PMN, indicating a change in transcriptome features at the final stages of the differentiation.

The P/M to MN and SN to PMN were the consecutive transitions with the highest number of differentially expressed genes and changes in chromatin states. These transcriptomic and epigenomic differences reflect the important differences between these stages. Indeed the P/M to MN denotes a passage from cells capable of dividing to cells with indented nuclei and unable to divide (Wahed and Dasgupta, 2015). Similarly, the SN to PMN transition denotes the passage from bone marrow to the bloodstream.

Different stages of maturation do not display relevant differences in DNA methylation. This suggests that the unique patterns of DNA methylation found in neutrophils (Schuyler et al., 2016) are established at earlier stages of differentiation, in agreement with a previous study (Ronnerblad et al., 2014) reporting that only minor changes occur in the P/M to PMN transition.

In line with the DNA methylation results the two repressive histone marks analyzed cannot be used to discriminate among neutrophil differentiation stages. Previous studies demonstrated that repressive H3K9me3 and H3K27me3 marks and DNA methylation have a crucial role in cell commitment (Bock et al., 2012, Hawkins et al., 2010) and our results suggest that, in neutrophils, they do not play any further roles once lineage commitment is achieved.

We focused our attention on promoter and enhancer regions subject to dynamic acetylation across differentiation. These regions, subdivided in clusters, showed significant overlaps of target genes with the clusters of differentially expressed genes, highlighting a strong correlation between epigenomic and transcriptomic profiles at the various differentiation stages. The study of dynamically acetylated regions led us to identify super-enhancers differently modulated across differentiation; a consistent group of these is activated at the end of differentiation and have as potential target the genes involved in immunological response and neutrophil activation.

To further characterize the dynamically acetylated regions we looked for enriched TF binding sites and integrated them with the differential expression results. This approach aimed to highlight TFs playing a potential role in the differentiation process. CEBPA is a TF of the bZIP family with a major role in the neutrophil differentiation program, and it is known to be crucial for the development of acute myeloid leukemia (Avellino and Delwel, 2017). Our results show that CEBPA is highly expressed in the bone marrow and then significantly down-regulated in the SN to PMN transition. CEBPE, another member of the bZIP family, is also highly expressed in the bone marrow and is down-regulated in the SN to PMN transition but also in the preceding transition BN to SN. Interestingly CEBPE modulates some of the lactoferrin-containing specific granule genes (Khanna-Gupta et al., 2007) and regulates the expression of GG and FG granule genes (Gombart et al., 2001), both characterized by a decreasing trend of expression at the PMN stage.

The transcriptome and epigenome characterizations depicted a precise differentiation program that, in agreement with previous findings (Theilgaard-Monch et al., 2005), progressively activates immune functionalities turning off essential cell processes. The study of five differentiation stages gave us the opportunity to detect with better resolution where the cell-cycle inactivation takes place. We have shown that the P/M stage is the most proliferative, with Ki67 signal dimming in the following MM and BN stages and to completely disappear in the SN stage. These data, alongside with the expression of key cell cycle regulators, indicate that the cell cycle is arrested after the P/M stage.

The antimicrobial activity of neutrophils is achieved through ROS production by the NADPH oxidase enzyme complex. We showed that this enzyme complex activity reaches its peak only at the end of differentiation, although its expression is already detectable at the MM stage. The observed difference in ROS production during differentiation between PMA and PAF/fMLP kinetics highlights that G-protein coupled receptor and its signaling machinery are required to trigger the process. Indeed, we showed that when the surface expression of fMLP-receptor FPR1 is low, at early stages of development, the response to PAF/fMLP is impaired.

Along the same line, we noticed that genes encoding for the proteins contained in the various granules displayed different transcriptional and epigenetic dynamics. Indeed the proteases were released by cellular activation only at the mature stage because the signaling cascade and surface expression of activating GPCR receptors were not present at earlier stages.

The results presented in this study shed light on the neutrophil differentiation processes and represent a useful resource for future studies on as yet uncharacterized primary immune disorders or neutrophil-related disorders like congenital neutropenia syndromes, Chediak Higashi syndrome and Lazy Leukocyte syndrome.

## Acknowledgements

The work presented here was supported by the Landsteiner Foundation for Blood Transfusion Research (LSBR-1121) to T.W.K. and by the Dutch Cancer Foundation (KUN 2011-4937) to J.A.H.M.. The work was funded by a grant from the European Commission 7th Framework Program (FP7/2007-2013, grant 282510, BLUEPRINT). M.F. is supported by the British Heart Foundation (BHF) Cambridge Centre of Excellence (RE/13/6/30180). Additional support was provided by the Wellcome Trust (WT108749/Z/15/Z) (P.F.) and by the European Molecular Biology Laboratory (E.L., A.D., L.C., P.F.). Research in the W.H.O. laboratory is also supported by grants from Bristol Myers-Squibb, British Heart Foundation, Medical Research Council, NIHR (W.H.O. is NIHR Senior Investigator) and NHS Blood and Transplant (NHSBT).

The web site https://blueprint.haem.cam.ac.uk/neutrodiff collects the analyzed data presented in this article. All the genome-wide data generated in this study, as well as the sequencing details, can be accessed via http://dcc.blueprint-epigenome.eu/a.

## Authorship Contributions

F.P. and T. W. K., designed the study, performed and supervised research, and wrote the manuscript; L.G. analyzed the data and wrote the manuscript; S.U., A.M., F.W, E.C., G.Y. and J.P. analyzed the data; I.H.H., A.T.J.T., E.M., E.J.M., K.B., H.K., E.H., M.G., M.L.Y. and M.L. performed experiments. E.L., A.D., L.C. and P.F. performed and supervised data management. M.V., D. Roos, T.K.vdB., S.H., D. Rico, M.K., I.G., A.V., W.H.O. and H.G.S. provided expert supervision. M.F. provided expert supervision and wrote the manuscript. J.H.A.M. designed the study, analyzed the data, supervised research and wrote the manuscript. All authors read and approved the final version of the manuscript.

## Experimental Procedures

### Study design

Bone marrow neutrophils have been collected from three different healthy donors and mature neutrophils have been collected from other three other different healthy donors. ChiP-Seq, RNA-Seq and WBGS sequencing experiments have been done on samples from the same donors.

### Cell isolation and library preparation

The study was approved by the local ethical committee of Sanquin Blood Supply Organization, and the Academic Medical Center, Amsterdam, The Netherlands. PMNs were enriched by gradient centrifugation and purified by negative selection with EasySep™ Human Neutrophil Enrichment Kit. Bone marrow neutrophil progenitors were purified by FACS with CD11b and CD16 (see supplemental methods for further details). ChIP-Seq and total RNA-Seq libraries were prepared according to the BLUEPRINT protocols (http://www.blueprint-epigenome.eu/).

### ChIP-Seq analysis and genome segmentation

Sequenced reads were mapped against the GRCh38 genome assembly with BWA (Li and Durbin, 2009). ChromHMM software (v1.10) (Ernst and Kellis, 2012) was then used to segment the genome in 200 bp intervals and assign epigenetic states. Chromatin state transitions consistently identified in all three replicates were represented with a Sankey diagram at each stage of development using the “makeRiver” and “riverplot” functions included in the riverplot R package v0.5.

### RNA-Seq analysis

Trim Galore (v0.3.7) (http://www.bioinformatics.babraham.ac.uk/projects/trim_galore/) with parameters “-q 15 -s 3 --length 30 -e 0.05” was used to trim PCR and sequencing adapters. Trimmed reads were aligned to the Ensembl v80 (Cunningham et al., 2015) human transcriptome with Bowtie 1.0.1 (Langmead et al., 2009) using the parameters “-a --best --strata -S -m 100 -X 500 --chunkmbs 256 --nofw --fr”. MMSEQ (v1.0.8a) (Turro et al., 2014, Turro et al., 2011) was used with default parameters to quantify gene expression. Genes with posterior probability > 0.5 (calculated by MMDIFF), absolute fold change > 2 and FPKM > 1 in at least one of the two compared cell types were considered differentially expressed. Differentially expressed genes in at least one of the comparisons P/M-MM, MM-BN, BN-SN, SN-PMN, P/M-PMN (supplemental Figure 1) were clustered using the “kmeans” (Hartigan and Wong, 1979) clustering R function. Gene expression FPKMs were log2 transformed and for each gene the z-score was calculated. We set the number of centers to seven according to the results of the gap statistic (Tibshirani et al., 2001) by using the “clusGap” function of the cluster R package.

### Super-enhancer identification

Super-enhancers (SEs) in each sample were predicted by the ROSE algorithm (Whyte et al., 2013) using H3K27ac as the surrogate mark. Briefly, all H3K27ac peaks within +/− 2.0□kb around transcription start sites (TSSs) were first excluded. The remaining peaks closer than default distance of 12.5 kb were stitched together, and subsequently ranked by normalized H3K27ac level corrected by input background. Finally, SEs were separated from typical enhancers based on the inflection point of H3K27ac signal curve.

### Functional enrichment analysis

Functional enrichment analysis was done by using FIDEA (D’Andrea et al., 2013) with default settings.

### Promoter/Enhancer/Super-enhancer gene assignment

For each transcript annotated in Ensembl v80 (Cunningham et al., 2015), we defined a promoter as the 2000 base pairs (bp) (± 1000 bp) around its transcription starting site. All the regions defined as promoter by the genome segmentation analysis and the H3K27ac dynamic analysis were assigned to the genes of the overlapping promoters. All the regions defined as enhancer by segmentation analysis and the H3K27ac analysis were assigned to the gene with the closest promoter (distant less than 10,000 bp) and, as well, to genes with promoters overlapping their interacting regions (CHiCAGO score >=5 in Neutrophils) derived from the Javierre *et al.* study (Javierre et al., 2016). The “findOverlaps” and “distanceToNearest” functions of the GenomicRanges (Lawrence et al., 2013) R package have been used to make region comparisons.

### Motif analysis

was performed with the HOMER (Heinz et al., 2010) program.

### WGBS sequencing

was carried out to minimum genome coverage of 30x with established protocols (Kulis et al., 2012).

### Differential methylation

was estimated for a) individual CpG sites in pairwise comparisons by using methyl_diff (Raineri et al., 2014) and across regions in group-wise comparison by employing replicates with metilene (Juhling et al., 2016).

### Association between acetylation and expression clusters

We built a contingency table with the number of genes for each acetylation cluster (rows) and gene expression cluster (columns). The Pearson’s chi-squared test was used to verify the non-independence of the two categorical variables. The significance of the association of each gene expression cluster with each dynamic acetylation cluster has been tested by a hyper-geometric test and the derived *P* values have been corrected with Benjamini & Hochberg method (Hochberg and Benjamini, 1990).

### NADPH-oxidase activity

was assessed as the release of hydrogen peroxide determined with an Amplex Red kit (Drewniak et al., 2013, Kuijpers et al., 2005).

### Protease release

after degranulation was measured with DQ-green BSA (see supplemental methods for further details).

